# When a foundation crumbles: forecasting forest community dynamics following the decline of the foundation species *Tsuga canadensis*

**DOI:** 10.1101/099333

**Authors:** Bradley S. Case, Hannah L. Buckley, Audrey A. Barker-Plotkin, David A. Orwig, Aaron M. Ellison

## Abstract

In the forests of northeastern North America, invasive insects and pathogens are causing major declines in some tree species and a subsequent reorganization of associated forest communities. Using observations and experiments to investigate the consequences of such declines are hampered because trees are long-lived. Simulation models can provide a means to forecast possible futures based on different scenarios of tree species decline, death, and removal. Such modeling is particularly urgent for species such as eastern hemlock (*Tsuga canadensis*), a foundation species in many northeast forest regions that is declining due to the hemlock woolly adelgid (*Adelges tsugae*). Here, we used an individual-based forest simulator, SORTIE-ND, to forecast changes in forest communities in central Massachusetts over the next 200 years under a range of scenarios: a no-adelgid, status-quo scenario; partial resistance of hemlock to the adelgid; adelgid irruption and total hemlock decline over 25 years, adelgid irruption and salvage logging of hemlock trees; and two scenarios of preemptive logging of hemlock and hemlock/white pine.We applied the model to six study plots comprising a range of initial species mixtures, abundances, and levels of hemlock dominance. Simulations indicated that eastern white pine, and to a lesser extent black birch and American beech, would gain most in relative abundance and basal area following hemlock decline. The relative dominance of these species depended on initial conditions and the amount of hemlock mortality, and their combined effect on neighborhood-scale community dynamics. Simulated outcomes were little different whether hemlock died out gradually due to the adelgid or disappeared rapidly following logging. However, if eastern hemlock were to become partially resistant to the adelgid, hemlock would be able to retain its dominance despite substantial losses of basal area. Our modeling highlights the complexities associated with secondary forest succession due to ongoing hemlock decline and loss. We emphasize the need both for a precautionary approach in deciding between management intervention or simply doing nothing in these declining hemlock forests, and for clear aims and understanding regarding desired community- and ecosystem-level outcomes.

## INTRODUCTION

A major challenge for ecologists is to study and forecast the dynamics of long-lived species, the assemblages they form, and the ecosystems they inhabit. These challenges are especially acute for studies of trees and forests, in which individuals can live for millennia and for which rates of change of some ecosystem processes occur over centuries. In many cases, researchers have used results from short-term observations and experiments to parameterize simulation models (Scheller and Mladenoff 2007). Subsequent experiments with model parameters inform research about, and suggest management strategies for, the emerging forests of the future (e.g., Cyr et al. 2009, Fontes et al. 2010). Such studies increasingly are of critical importance because many forests around the world are in decline (Butchart et al. 2010) and are changing rapidly (Choat et al. 2012, Lindenmayer et al. 2012).

Perhaps nowhere are these changes more apparent than in northeastern North America, where a number of tree species are declining due to irruptions of native and nonnative insects and pathogens (Dukes et al. 2009). Of particular note is the rapid decline and disappearance of the forest foundation species (*sensu* Ellison et al. 2005), eastern hemlock (*Tsuga canadensis* (L.) Carr.), resulting from spread of a nonnative insect, the hemlock woolly adelgid (*Adelges tsugae* Annand), along with the removal of this tree from forests by pre-emptive salvage logging (Orwig and Kittredge 2005, Foster and Orwig 2006, Fajvan 2007, Ellison et al. 2010). Many researchers have dedicated substantial time and effort in quantifying the pattern and rate of spread of the adelgid (Preisser et al. 2008, Morin et al. 2009, Fitzpatrick et al. 2010, 2012, Gomez et al. 2015), the rate and intensity of adelgid-related mortality for different demographic stages of hemlock (Preisser et al. 2011), and the ultimate effects of hemlock mortality on forest ecosystem properties and dynamics (Templer and McCann 2010, Block et al. 2012, Orwig et al. 2013).

Eastern hemlock is a long-lived, late-successional conifer species and, where it occurs, often comprises a large proportion of overstorey basal area, either as pure stands, or as the dominant species in mixed stands with other species such as eastern white pine (*Pinus strobus* L.), red maple (*Acer rubrum* L.), birch species (*Betula* spp.), and oaks (*Quercus* spp.) (Orwig et al. 2002). The loss of eastern hemlock from these forests results in the formation of new, multi-sized canopy gaps, causing an associated increase of light into the heavily-shaded and generally species-poor understory layer typical of hemlock stands (Ellison et al. 2010). Possible successional pathways in response to this situation will be contingent largely on gap size, overall site conditions, and the proximity, population structure, and life history attributes of the co-occurring species in the community (Small et al. 2005, Eschtruth et al. 2006, Ellison et al. 2010, Ford et al. 2012). Experimental (Ellison 2014) and observational studies (Small et al. 2005, Eschtruth et al. 2013) have provided insights into possible trajectories of future forest development following the loss of eastern hemlock, but these studies are of relatively short duration (< 20 years). Process-based simulation models provide a means to explore a range of scenarios associated with insect irruptions and subsequent changes in forest structure and composition over a range of spatiotemporal extents and resolutions (Papaik et al. 2005, Cairns et al. 2008).

Very few modeling studies have explored how forest ecosystem dynamics will change following loss of eastern hemlock (Albani et al. 2010, Birt et al. 2014). Individual-based modeling approaches should be particularly useful in exploring how tree- and neighborhood-scale dynamics influence trajectories of species composition and structure under hemlock decline scenarios that are relevant to a specific forest area (Jenkins et al. 2000). Here we use an individual-based forest simulator, SORTIE-ND (Pacala et al. 1993, 1996, Canham et al. 2004, 2006), parameterized for northeastern North American forests and initialized with tree-level data from plots within the Harvard Forest in central Massachusetts, to model how forest stands currently dominated by eastern hemlock may reorganize over the next 200 years under a range of scenarios of hemlock woolly adelgid-related impacts. SORTIE-ND incorporates plant demography, inter- and intraspecific interactions, and the effects of both natural and anthropogenic disturbance in its forecasts of future forest structure. Our models address three key questions: 1) what is the effect of adelgid-induced mortality on the next 200 years on structure and species composition of New England (USA) forests; (2) how different will these impacts be if partial resistance to the adelgid emerges in the next several decades; and (3) what are the interactions between adelgid-induced mortality, pre-emptive salvage logging of eastern hemlock by people, and background (“natural”) rates of hemlock mortality?

## METHODS

### Study area

We modeled the future of forest stands in the “Transition Hardwoods-White Pine-Hemlock” region (Westveld et al. 1956) of central Massachusetts. In many areas in northeastern North America, including our study sites at Harvard Forest, eastern hemlock may account for > 70% of the basal area of the trees within any given forest stand (Ellison et al. 2010, Orwig et al. 2012). The forests in central Massachusetts are underlain by loamy soils derived from acidic, glacial till; average annual temperatures are 7.1 °C; and average annual precipitation ≈1,000 mm (Greenland and Kittel 1997). Historically, much of the region was cleared for agriculture or harvested for timber in the early-to-mid 1800s, and has been regenerating since then (Foster 1992).

The hemlock woolly adelgid arrived in Massachusetts in the first decade of the 21^st^ century (Preisser et al. 2008). The adelgid was introduced from Asia into eastern North America (near Richmond, Virginia) in the early 1950s (Havill and Montgomery 2008). Since then, it has spread rapidly and has established itself in forests from Georgia northward into Canada and west into Michigan (Morin et al. 2009). The hemlock woolly adelgid in North America reproduces asexually and kills trees of all sizes, ages, and stages (Orwig and Foster 1998). Tree morbidity can occur within two years after adelgid establishment, and tree mortality follows in the next few to 15 years (Orwig et al. 2002). Further, in anticipation of the arrival of the adelgid in a region, people often log existing stands of eastern hemlock (Foster and Orwig 2006, Ellison et al. 2010). Goals of this logging include: increasing the vigor and potential survivorship of remaining eastern hemlock trees; redirecting succession and influencing future forest dynamics; and realizing economic return from trees that will otherwise die (Orwig and Kittredge 2005, Fajvan 2007). Because eastern hemlock is of low commercial value, people often harvest other more valuable species such as eastern white pine or red oak (*Quercus rubra* L.) at the same time as they log eastern hemlock (Brooks 2004). Logging and the adelgid have different short-term effects on forest succession and ecosystem dynamics because of differences rates of mortality and biomass removal (fast *versus* slow) and subsequent dynamics (Kizlinski et al. 2002, Orwig et al. 2013).

### SORTIE core parameterization

SORTIE-ND is a spatially explicit forest simulator that includes effects of tree neighbors on species-specific sapling and adult tree growth and mortality (Pacala et al. 1993, 1996, Canham et al. 2004, Canham et al. 2006). SORTIE-ND tracks the fates of individual trees located within a forest plot as the trees compete with neighboring trees for light and space. The basic setup of the model requires parameterization of equations related to the generation of light conditions across a given forest area, seed dispersal and seedling recruitment, and the growth, mortality, senescence, and death of all seedlings, saplings and adults of each species (Table 1). These equations include variables that characterize particular aspects of the life histories of individual species, including how they respond to light conditions and the crowding and shading effects of neighboring trees (Canham et al. 2006, Uriarte et al. 2009).

**Table 1.**
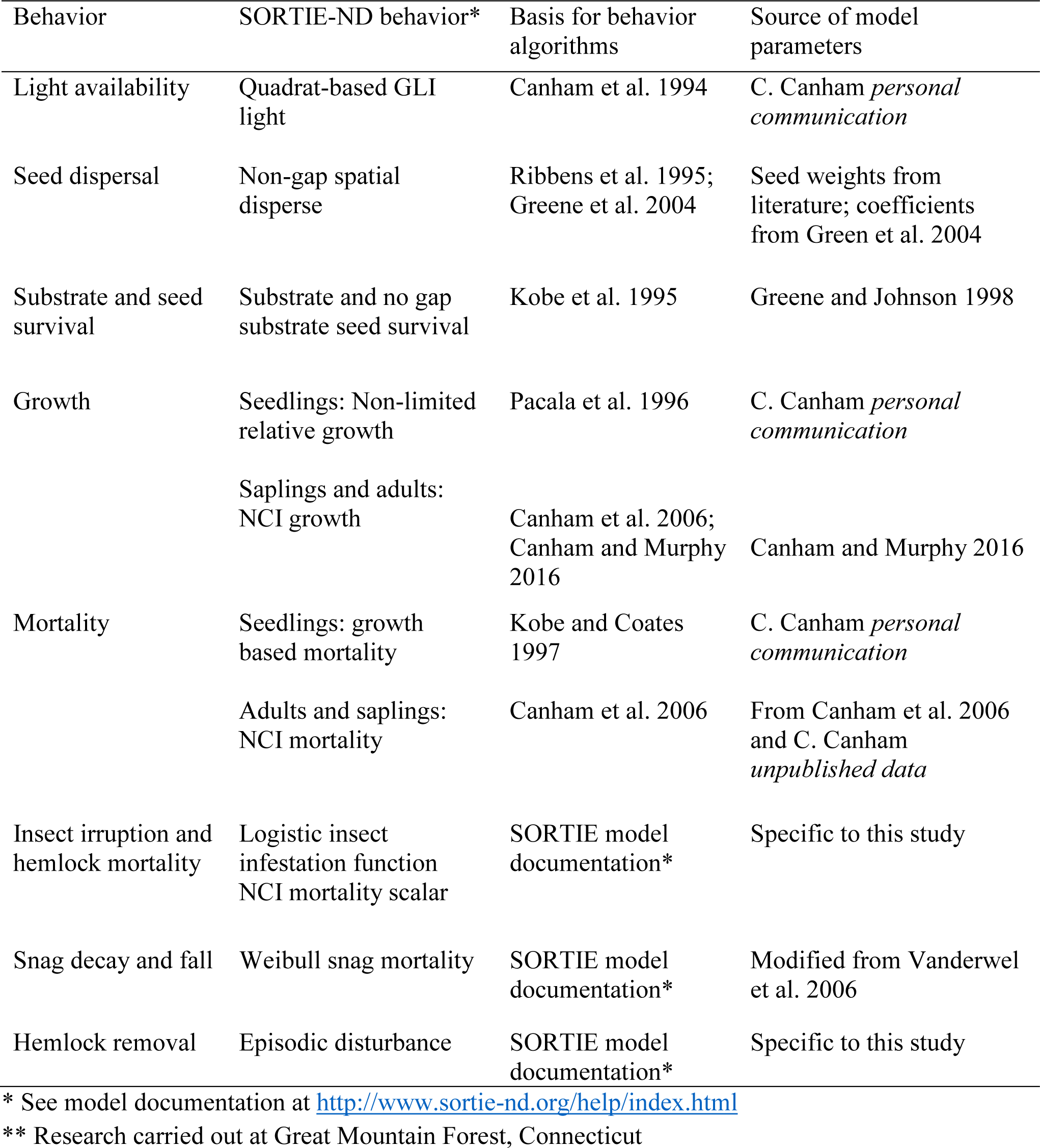
Overview of the main SORTIE-ND behaviors and parameters used for simulations.

| Behavior | SORTIE-ND behavior* | Basis for behavior algorithms | Source of model parameters |
| --- | --- | --- | --- |
| Light availability | Quadrat-based GLI light | Canham et al. 1994 | C. Canham *personal communication* |
| Seed dispersal | Non-gap spatial disperse | Ribbens et al. 1995; Greene et al. 2004 | Seed weights from literature; coefficients from Green et al. 2004 |
| Substrate and seed survival | Substrate and no gap substrate seed survival | Kobe et al. 1995 | Greene and Johnson 1998 |
| Growth | Seedlings: Non-limited relative growth | Pacala et al. 1996 | C. Canham *personal communication* |
|  | Saplings and adults: NCI growth | Canham et al. 2006; Canham and Murphy 2016 | Canham and Murphy 2016 |
| Mortality | Seedlings: growth based mortality | Kobe and Coates 1997 | C. Canham *personal communication* |
|  | Adults and saplings: NCI mortality | Canham et al. 2006 | From Canham et al. 2006 and C. Canham *unpublished data* |
| Insect irruption and hemlock mortality | Logistic insect infestation function<br>NCI mortality scalar | SORTIE model documentation* | Specific to this study |
| Snag decay and fall | Weibull snag mortality | SORTIE model documentation* | Modified from Vanderwel et al. 2006 |
| Hemlock removal | Episodic disturbance | SORTIE model documentation* | Specific to this study |

\* See model documentation at http://www.sortie-nd.org/help/index.html
\*\* Research carried out at Great Mountain Forest, Connecticut

The functions that determine recruitment of tree seedlings in SORTIE-ND are derived from field data describing seed production, dispersal, and establishment in a variety of soils (Ribbens et al. 1994, LePage et al. 2000, Greene et al. 2004, Papaik et al. 2005, Papaik and Canham 2006). In our implementation of SORTIE-ND, we used the “spatial disperse” option that disperses seeds in each time step at random azimuth directions and distances away from each tree of reproductive age or older. Dispersal distances follow a Weibull distribution, which results in an inverse relationship between the number of dispersed seeds and distance from the parent tree. Maximum fecundity for each species is based on the “standardized total recruits”: the total number of seeds produced by a 30-cm diameter parent tree, estimated as the average number of viable seeds kg^−1^ of seed species^−1^ (Ribbens et al. 1994). Seeds can fall on six different types of substrate: forest floor litter, moss, scarified soil, tip-up mounds, decayed logs, or fresh logs. The distribution and availability of each of these substrates are represented as a regular 2 × 2-m grid across the simulation area, and are based on the number of dead trees that fall each time step, the direction of tree fall, decay rate of fallen dead trees, and proportion of fallen trees that uproot and make tip-up mounds (Papaik et al. 2005). Successful seedling establishment in each grid cell was determined by species-specific affinity for a particular substrate and relative seed mass (Greene and Johnson 1998).

Seedling growth in SORTIE-ND uses a Michaelis-Menton function that increments annual diameter growth as a species-specific function of light availability (Pacala et al. 1996), computed within the 2 × 2-m grid cells. Available light is computed as a “global light index” (GLI), an estimate of whole-season photosynthetically-active radiation conditional on the crown geometry of individuals of each species, the location, size, and identity of nearby trees, and species-specific light transmission coefficients (Canham 1988, Canham et al. 1994, 1999). Species-specific seedling mortality was modeled as a function of the recent growth of each seeding, its shade tolerance, and within-cell light availability (Kobe et al. 1995). We used seedling growth and mortality parameters provided by C. Canham (*personal communication*) based on previous studies of New England forests (e.g., Pacala et al. 1993, Pacala et al. 1996). The growth and mortality of saplings and adults were modeled using neighborhood competition index (NCI) functions (Canham et al. 2004, 2006, Canham and Murphy 2016). The NCI functions adjust the maximum potential growth rate of each species by initial tree size, intra- and interspecific competition, and within-cell environmental conditions. We used parameters for sapling growth and mortality estimated as part of an analysis of Forest Service Forest Inventory and Analysis (FIA) data, investigating the effects of climatic change on forests across the eastern USA by Canham and Murphy (2016) and Canham (unpublished data).

### Model evaluation simulations

To evaluate SORTIE’s performance, we used 10 years of field observations on four plots within the Harvard Forest Hemlock Removal Experiment (HF-HeRE, see Ellison et al. 2010 and 2014 for details), located in the Simes Tract of the Harvard Forest Long Term Experimental Research (LTER) site (42.5°N, 72.2°W; mean elevation 250-m.a.s.l.; Fig. 1B). The HF-HeRE plots were established in 2003 to study experimentally the effects of adelgid-related hemlock mortality and preemptive salvage logging on forest structure and dynamics; post-treatment measurements of tree seedlings, saplings, and trees were done in 2009 and 2014. Specifically, we used the hemlock control plots numbers 3 and 6 (hereafter, “HeRE3-hemlock” and “HeRE6-hemlock”) and the two experimentally logged plots, plots 2 and 4 (hereafter, “HeRE2-logged” and “HeRE4-logged”). For the logged plots, we compared observed and simulated changes that had occurred after the 2005 logging operation. The application of SORTIE to the former two plots enabled us to evaluate how well the model can reproduce observed forest dynamics under a scenario of typical forest development without disturbance; the application to the latter two provided an evaluation of the model under a scenario of post-preemptive-logging of hemlock that has ostensibly occurred in advance of a catastrophic adelgid outbreak.

**Figure 1.**
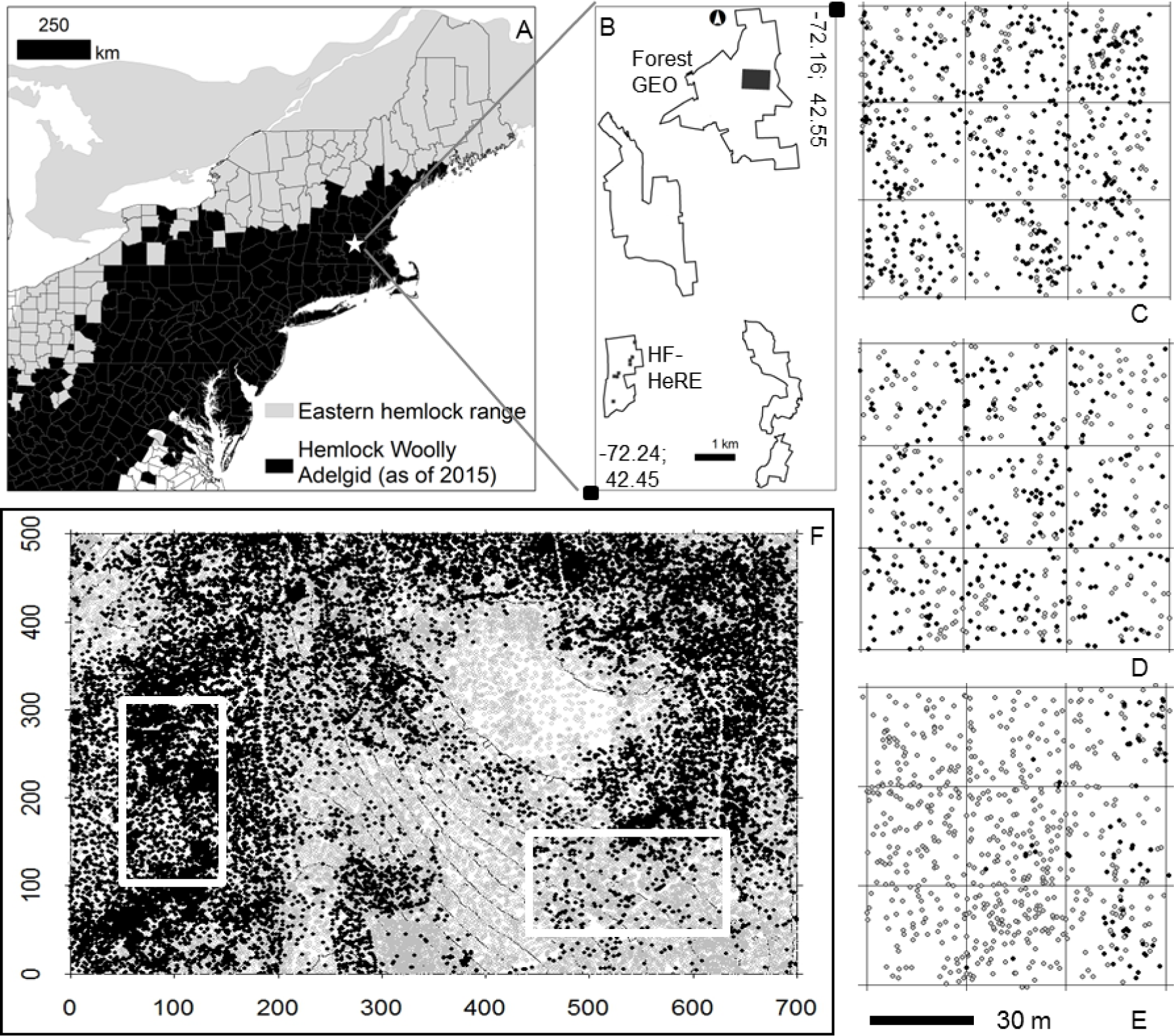
Study location in eastern North America, and stem maps of the plots used to initialize the forecasting simulation models. For the stem maps, black points are eastern hemlock and gray points are all other species. Clockwise from top left: A) range of eastern hemlock and counties with established hemlock woolly adelgid populations as of 2015 (data source: USFS, Northeastern Area; Disclaimer: This map depicts counties with established HWA populations that are confirmed and reported by respective state forest health officials. The coarse nature of the map does not provide information below the county level and readers should not assume that highlighted infested counties are entirely infested.); B) location of study plots at the Harvard Forest Long Term Experimental Research site in central Massachusetts, USA; C) HF-HeRE plot 3 (hemlock); D) HF-HeRE plot 6 (hemlock); E) HF-HeRE plot 7 (hardwood); F) ForestGEO plot with the two, 2 hectare sub-plots used indicated by the white outlines.

### Model forecasting simulations

We simulated forest dynamics for 200 years under six scenarios aimed at answering our main questions (Table 2). The first scenario, using the core parameterization of SORTIE described above, included only background levels of natural mortality of eastern hemlock. Scenarios 2 to 6 involved a range of hemlock mortality situations, via either adelgid-induced mortality (Scenarios 2 and 3), salvage logging (Scenario 4), preemptive logging of hemlock (Scenario 5), or logging of hemlock and merchantable pine (Scenario 6). These latter five scenarios required additional model parameterization.

**Table 2.**
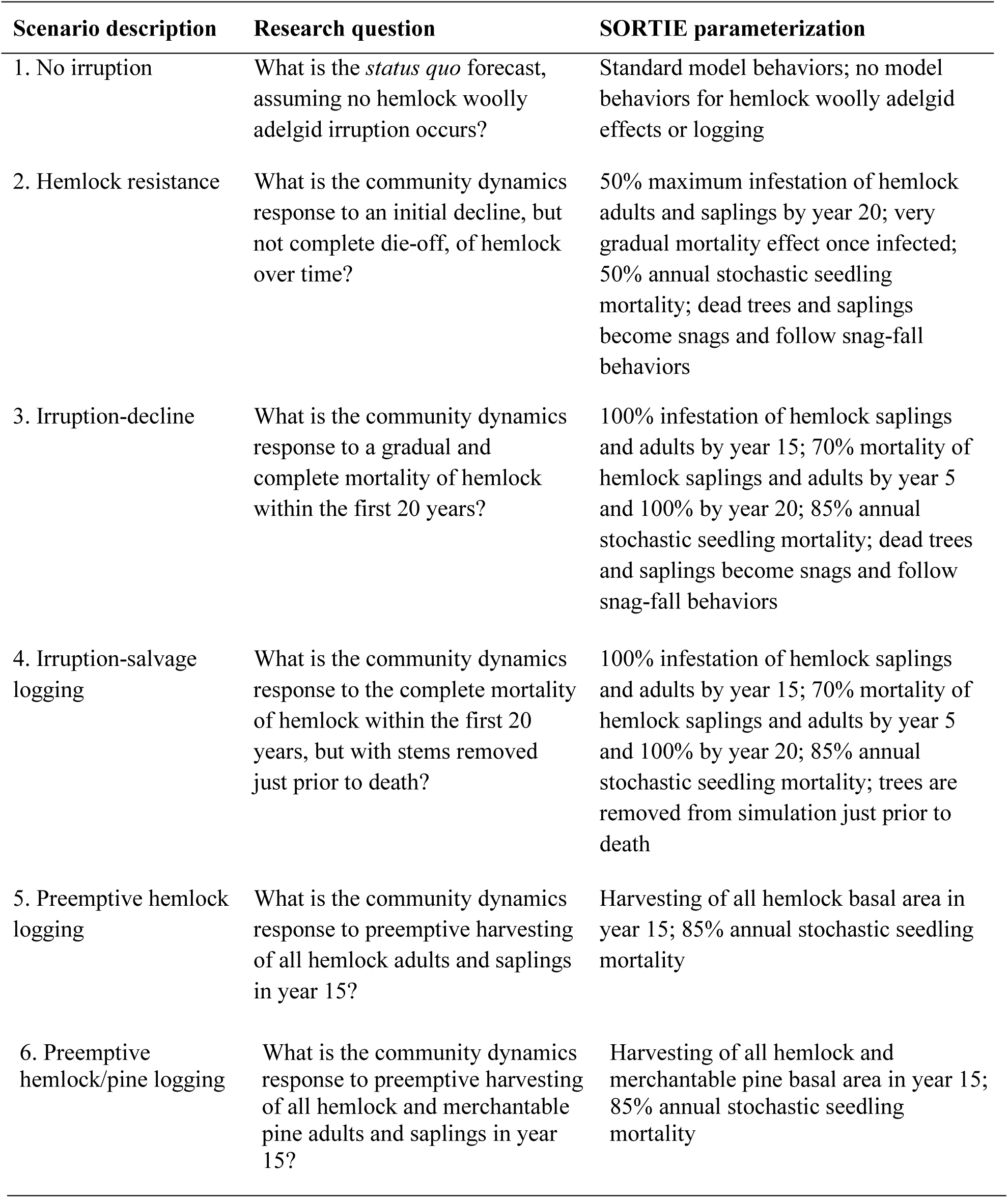
A description of the SORTIE-ND forecast simulations carried out at the Harvard Forest ForestGEO and Hemlock Removal Experiment plots. The proposed scenarios investigate possible forest structure and composition changes associated with hemlock woolly adelgid (hemlock woolly adelgid)-related impacts.

For initial inputs to the model under all six scenarios, and to encompass a range of initial conditions typical of forests of this region, we used data from a total of six study plots (Fig. 1). The first three plots were from the aforementioned HF Hemlock Removal Experiment, including the two HF-HeRE hemlock control plots used for model evaluation (HeRE3-hemlock and HeRE6-hemlock) and one hardwood control plot (hereafter, “HeRE7-hardwood”). These plots are 0.76-ha, 0.81-ha, and 0.81-ha in size, respectively. The other three study plots were associated with the 35-ha Harvard Forest ForestGEO forest dynamics plot located in the Prospect Hill tract (Orwig et al. 2015), also at the Harvard Forest LTER. The ForestGEO plot is one of a network of large forest plots established as part of the Smithsonian Tropical Research Institute’s Center for Tropical Forest Science – Forest Global Earth Observatory (CTFS- ForestGEO) network of plots. For our modeling, we used data from the full ForestGEO plot (hereafter, “FG-full”) and two, 2-ha sub-regions of the plot that currently are dominated either by eastern hemlock (“FG-hemlock”) or red oak (“FG-hardwood”).

In all simulations, we initialized the models using data for the ten tree species that comprised > 95% of the total basal area of the plots (Table 3). Initial model input data for all plots comprised species identities, sizes (DBH) and *x-y* locations (tree maps) of field-measured trees. For the HF-HeRE plots, all trees were measured and located for individuals > 5 cm DBH and 1.3 m in height; saplings (DBH < 5 cm and height < 1.3 m) were counted but not located; at the HF-ForestGEO plots, all trees and saplings with DBH > 1-cm and height > 1.3-m were measured and located. For the HF-HeRE plots, we used means of ten years of seedling plot data as initial inputs (Table 4), while for the ForestGEO plots, we used a common seedling density of 250 seedlings per hectare for all species because seedling data were unavailable. Raw data for all plots are available online from the Harvard Forest Data Archive (http://harvardforest.fas.harvard.edu/harvard-forest-data-archive): datasets HF126 (HF-HeRE overstory vegetation), HF106 (HF-HeRE understory vegetation), and HF253 (HF ForestGEO data).

**Table 3.**
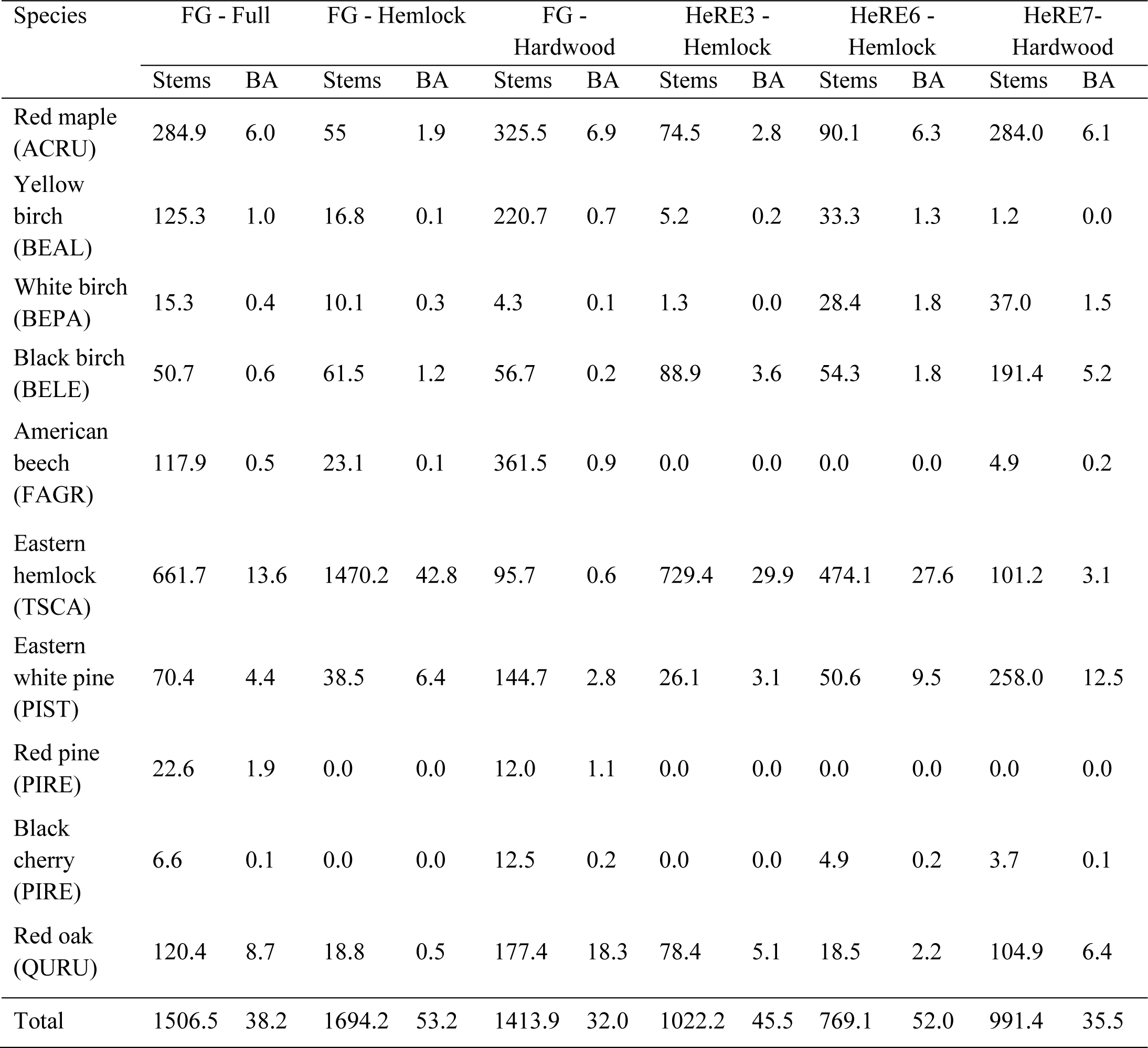
Observed adult and sapling stem densities (stem ha^−1^) and basal areas (m^2^ ha^−1^) for species (acronyms in parentheses) within the Harvard Forest ForestGeo (FG) and the Hemlock Removal Experiment (HeRE) study plots that comprise at least 90% of the total basal area (BA), and that were used as initial conditions for scenario simulations.

**Table 4.**
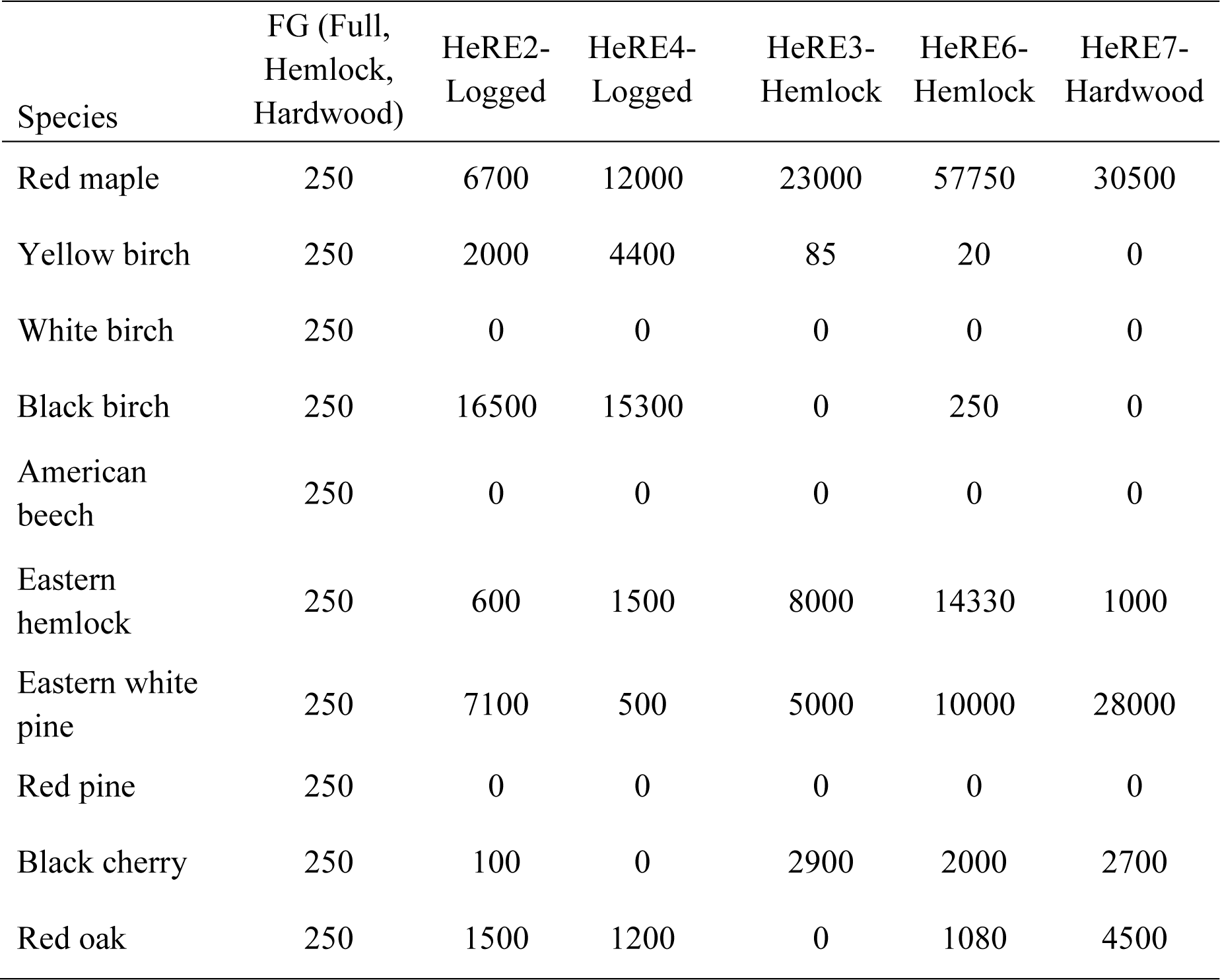
Initial seedling densities (seedlings ha^−1^) for model evaluation and scenario simulations for the Harvard Forest ForestGEO (FG) and Hemlock Removal Experiment (HeRE) study plots. A constant seedling density was used for the FG plots due to a lack of seedling data. Seedling values for the HeRE plots are means of seedling counts within 10, 1-m^2^ sample quadrats within each of the plots sampled over 10 years, scaled to a per-hectare basis.

To simulate adelgid-induced mortality in SORTIE-ND, we used a logistic equation (Table 1) that estimates the proportion of trees colonized by the adelgid. This equation includes parameters for initial and maximum rates of establishment, the steepness of establishment rate, and the time at which half the maximum rate of establishment has occurred. For hemlock trees, the NCI mortality function was adjusted with an additional parameter that changed the probability of mortality as a function of time that the adelgid had been present. In our simulations, we used two alternative parameter sets: one based on the scenario in which there was partial resistance of eastern hemlock to the adelgid (Scenario 2) and one in which there was no resistance of eastern hemlock, the adelgid spread rapidly, and all trees were killed by simulated year 25 (Scenario 3). We also included functions to allow the formation of standing dead trees (“snags”) and their subsequent fall (Venderwel et al. 2006). To mimic salvage logging (Scenario 4), we simply eliminated hemlock snag formation from the model, which forced the model to delete all hemlock individuals as they died. An episodic mortality behavior was used to simulate preemptive logging (Scenarios 5 and 6). Trees were killed and removed as a function of parameters that controlled the timing and spatial extent of each logging event, the range of diameters within which mortality occurred, and the proportion of total basal area within the specified diameter range that was removed. We affected complete logging of eastern hemlock (Scenario 5) or eastern hemlock and merchantable white pine (Scenario 6) at simulation year 15.

Stochastic behavior in SORTIE-ND results from differences in initial conditions and from using random draws from specified distributions for dispersal, growth, and mortality parameters. We used field data from six plots as our initial conditions and did not vary these among runs. However, comparisons of simulations initialized by different plots did allow us to examine how variation in initial community structure could result in different outcomes. We did use random draws from probability distributions for dispersal, growth, and mortality parameters. For each plot × scenario combination, we ran 10 stochastic simulations.

## RESULTS

### Model evaluation

Observed adult tree stem densities and basal areas changed from the starting conditions (2003-2004) to the first (2009) and second (2014) re-measurements in the two HF-HeRE hemlock control plots (plots 3 and 6) and two logged plots (plots 2 and 4) used for model evaluation (Figs. 2 and 3). Over the 10-year measurement period, stem densities decreased by 17.5% and 5.8% in plots 3 and 6, respectively, and increased by 2.9% and 17.1% post-logging in plots 2 and 4, respectively. On average, total plot basal area increased by 6.1% in plots 3 and 6 and by 24.6% post-logging in plots 2 and 4 over the same 10-year period. These observed changes in the distributions of stem densities (Fig. 2) and basal areas (Fig. 3) by diameter classes were well-simulated by SORTIE for both re-measurement times for plots 3, 6 and 2, but were under-predicted for most diameter classes of trees in plot 4.

**Figure 2.**
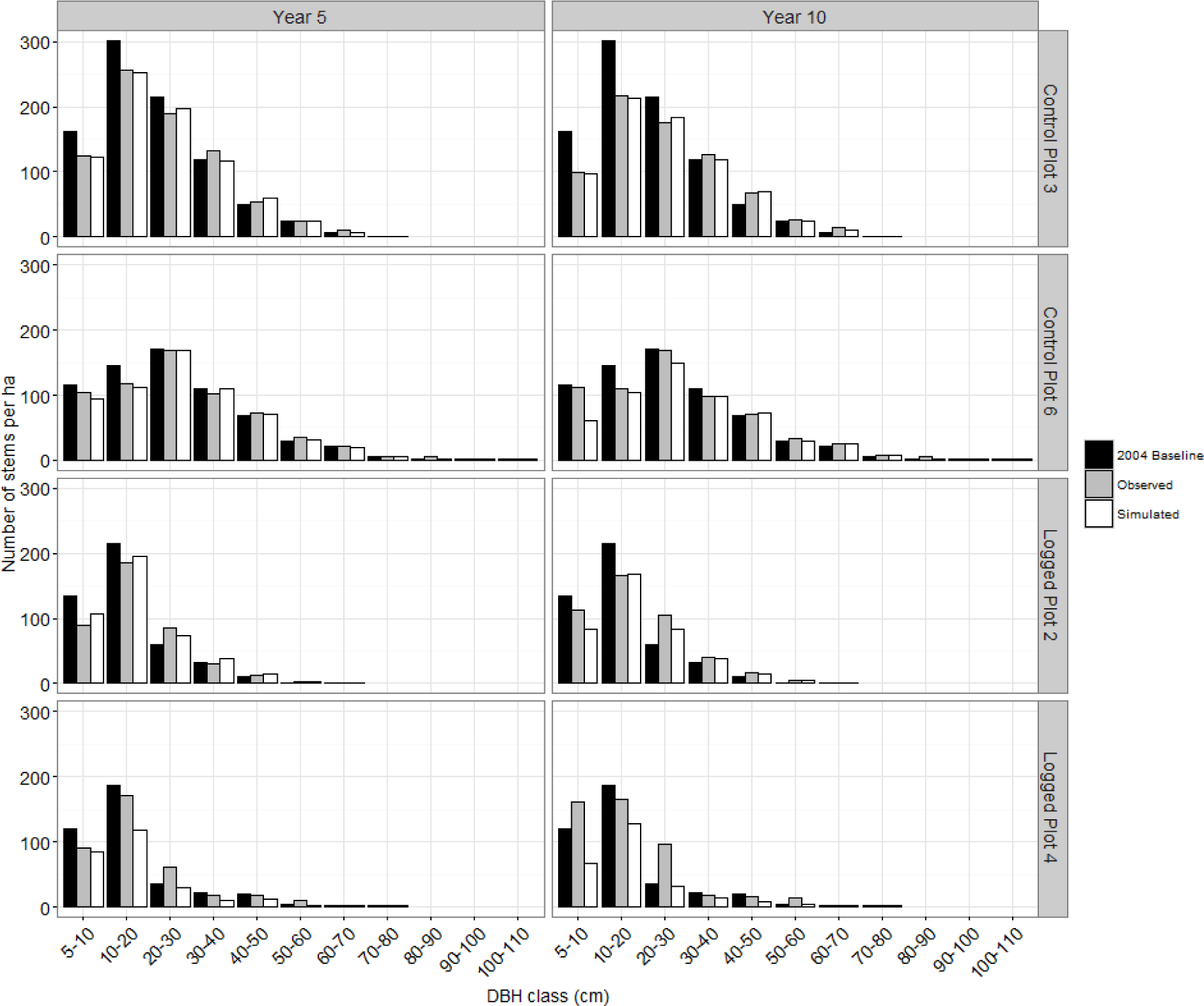
Comparison of observed and simulated live stem density distributions (A and B) and basal area distributions (C and D) by tree diameter categories after five years (first re-measurement, 2009) and ten years (second re-measurement, 2014) in the Harvard Forest Hemlock Removal Experiment hemlock control plots 3 and 6 and logged plots 2 and 4. For comparison purposes, we provide observed stem densities for the initial measurement (2004 Baseline).

**Figure 3.**
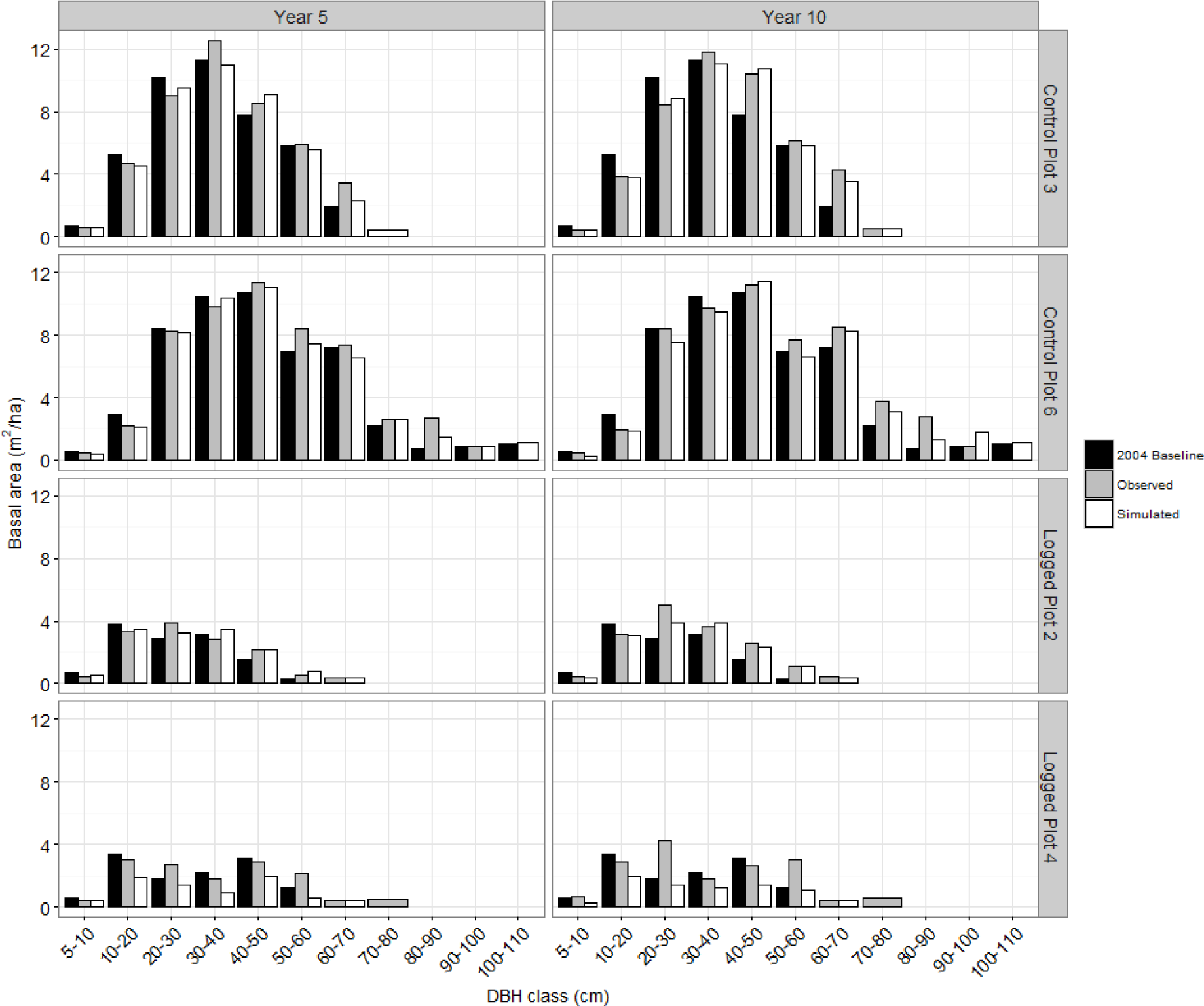
Comparison of observed and simulated basal area distributions by tree diameter categories after five years (first re-measurement, 2009) and ten years (second re-measurement, 2014) in the Harvard Forest Hemlock Removal Experiment for the hemlock control plots (3 and 6) and logged plots (2 and 4). For comparison purposes, we provide observed basal areas for the initial measurement (2004 Baseline).

After logging, observed black birch (*Betula lenta* L.) sapling densities increased dramatically in plots 2 and 4 after year four, from zero stems per hectare to near 10,000 stems per hectare in year 10 (Fig. 4). Saplings of other species were far less prominent in the plots, although there was a slight trend of increasing red maple saplings from year six and white pine from year eight. Similar to observed patterns, SORTIE also simulated a large influx of black birch saplings to over 10,000 stems per hectare, although this did not occur until after year nine (Fig. 4), indicating a lag in the movement of seedlings to the sapling tier in the model. The model similarly predicted an increase in red maple after year 14 in both plots and an increase in white pine in plot 4, while other species comprised relatively minor components. Model simulations indicated that over the next 20 years, although decreasing after its initial burst, black birch saplings will continue to dominate the gaps created by logging, with considerable numbers of red maple and, in plot 4, white pine persisting.

**Figure 4.**
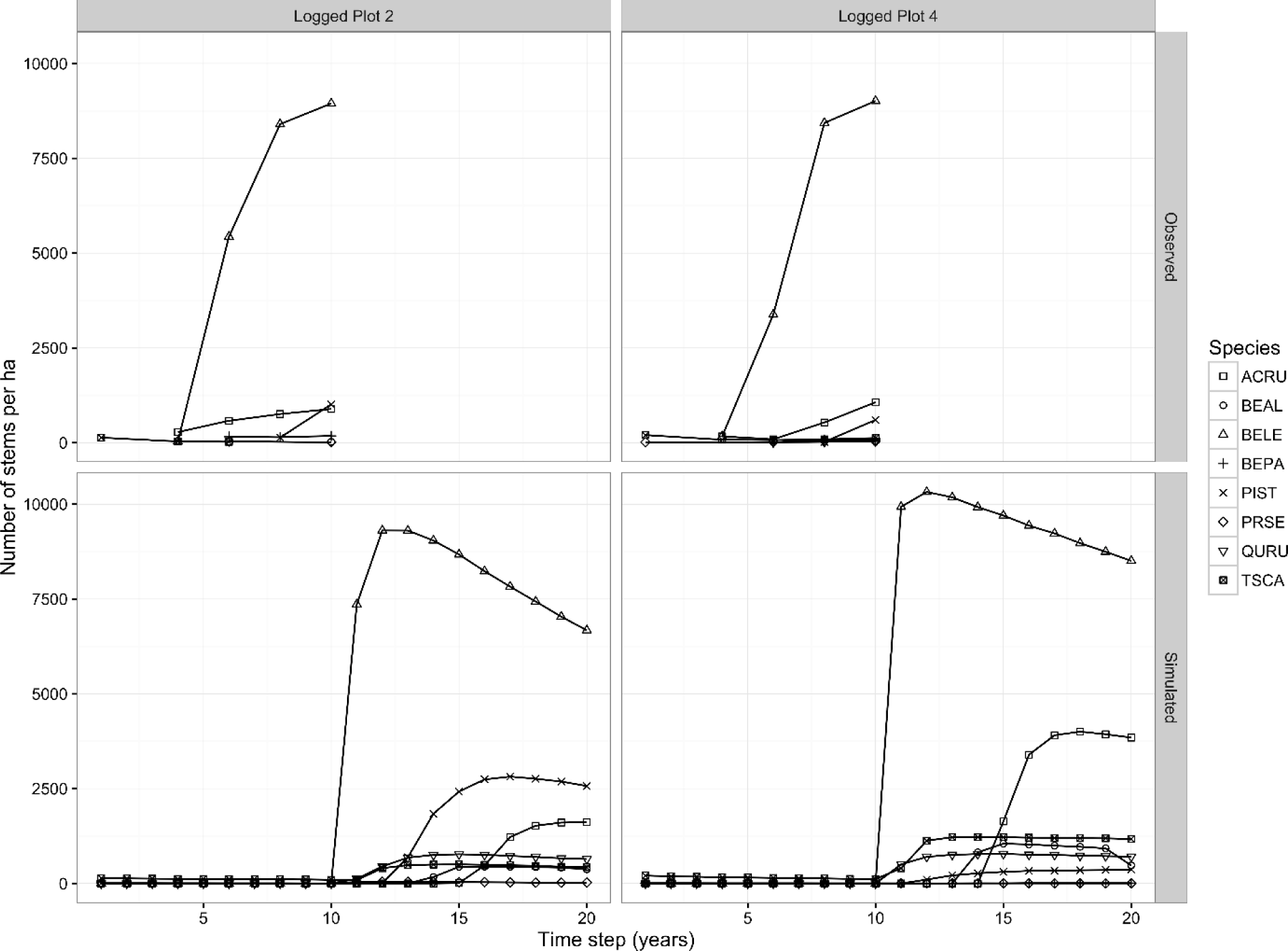
Comparison of observed and simulated live sapling density distributions in the Harvard Forest Hemlock Removal Experiment, Hemlock Logged Plots 2 and 4, by species.

### Forecasts – forest dynamics and structure

Under a no-adelgid-irruption scenario (Scenario 1), simulations suggested that in the four hemlock-dominant plots, hemlock would continue to self-recruit and maintain its dominance over time (Fig. 5); by the end of the simulation horizon, these dynamics resulted in an overall loss of total basal area due to higher numbers of hemlock individuals in smaller age classes (Figure 6). In the two hardwood-dominant plots, hemlock would also increase in prevalence as a canopy species, mainly replacing oak as it matures and dies over time (Fig. 5), increasing the overall basal area of these plots by year 200 (Fig. 6).

**Figure 5.**
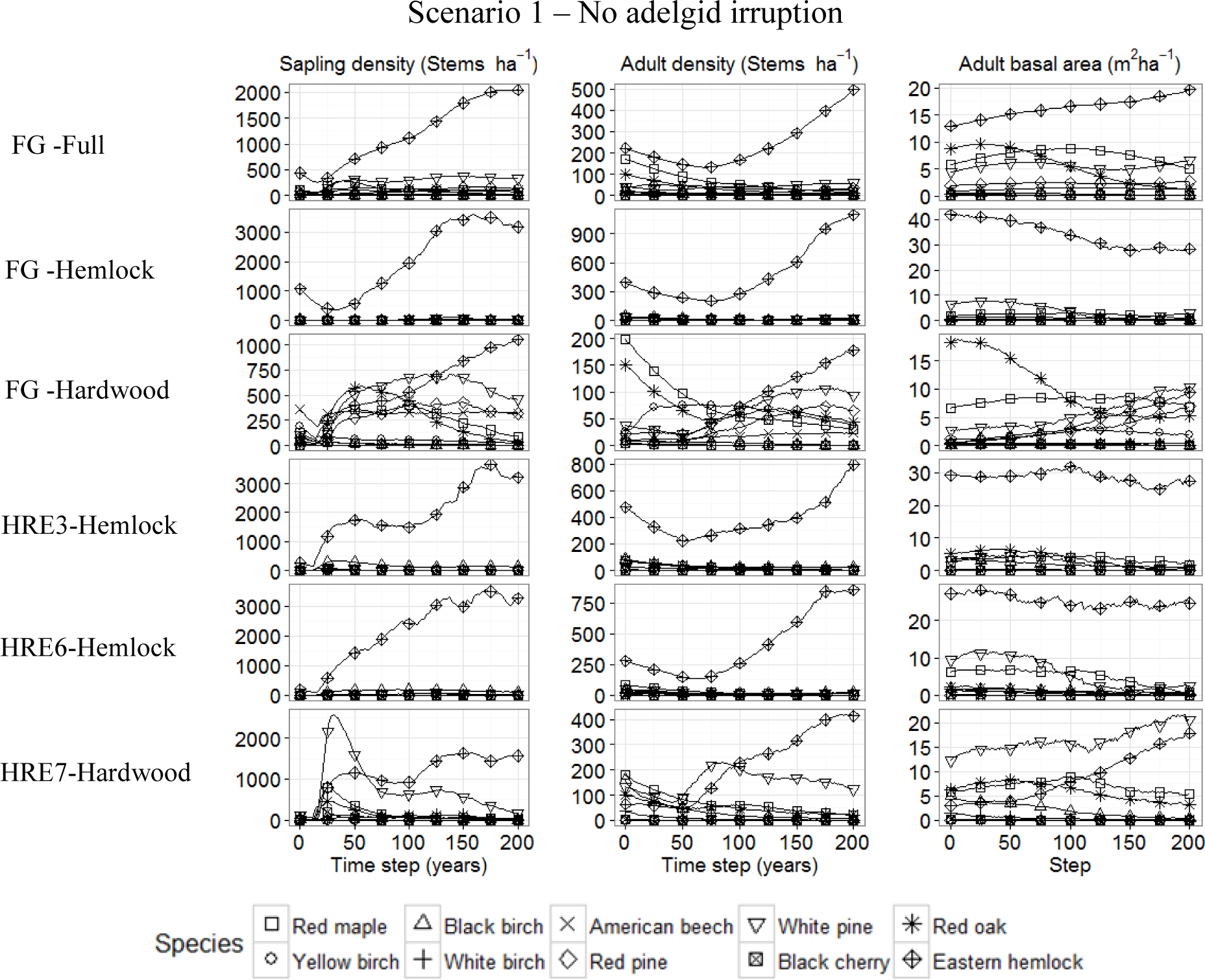
Simulated forest community dynamics over 200 years for Scenario 1 within the six study plots, characterized in terms of changes in sapling and adult stem density (stems ha^−1^) and adult basal area (m^2^ ha^−1^) for each of ten dominant tree species. Lines represent median values for each species of ten model simulations.

**Figure 6.**
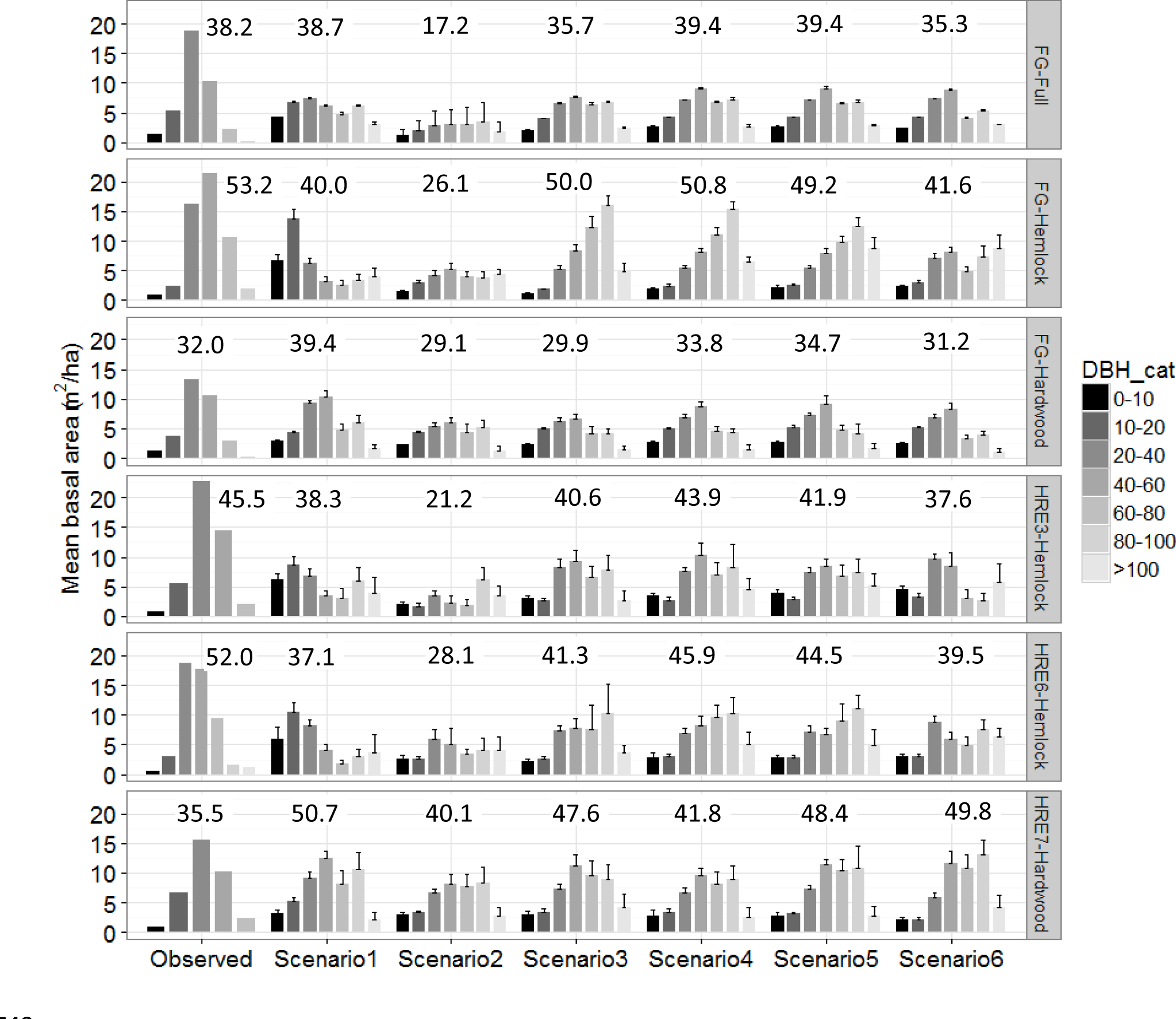
Basal area (m^2^ ha^−1^) distribution by dbh class for the six study plots for current (observed) forest conditions and for simulated, year 200 conditions under the six simulation scenarios (see Table 2). For observed data, values are total basal areas per dbh class; for simulated data, values are means of total basal area per dbh class across 10 model runs, with variation around the means indicated by two standard error bars. Above each group of bar graphs are total plot basal areas.

Under a scenario of partial hemlock resistance to adelgid impacts (Scenario 2), SORTIE simulated a variety of impacts on species composition and structure that depended largely on the initial composition of the plot. In the hemlock-dominant plots, American beech and black birch recruited as saplings to high densities into the gaps created by dying hemlock trees (Fig. 7), while hemlock concomitantly retained its overall dominance and eventually began to recover over time. The effect of this is that by year 200, there was an overall dampening of the size structure of these plots and a reduction of up to one-half the original total basal area (Fig. 6). By contrast, in the two hardwood-dominated plots, existing adult white pine and yellow birch also took advantage of these gaps, increasing their dominance in the canopy over time, resulting in only minor changes in the size structure of these plots relative to starting profiles (Figs. 7 and 6).

**Figure 7.**
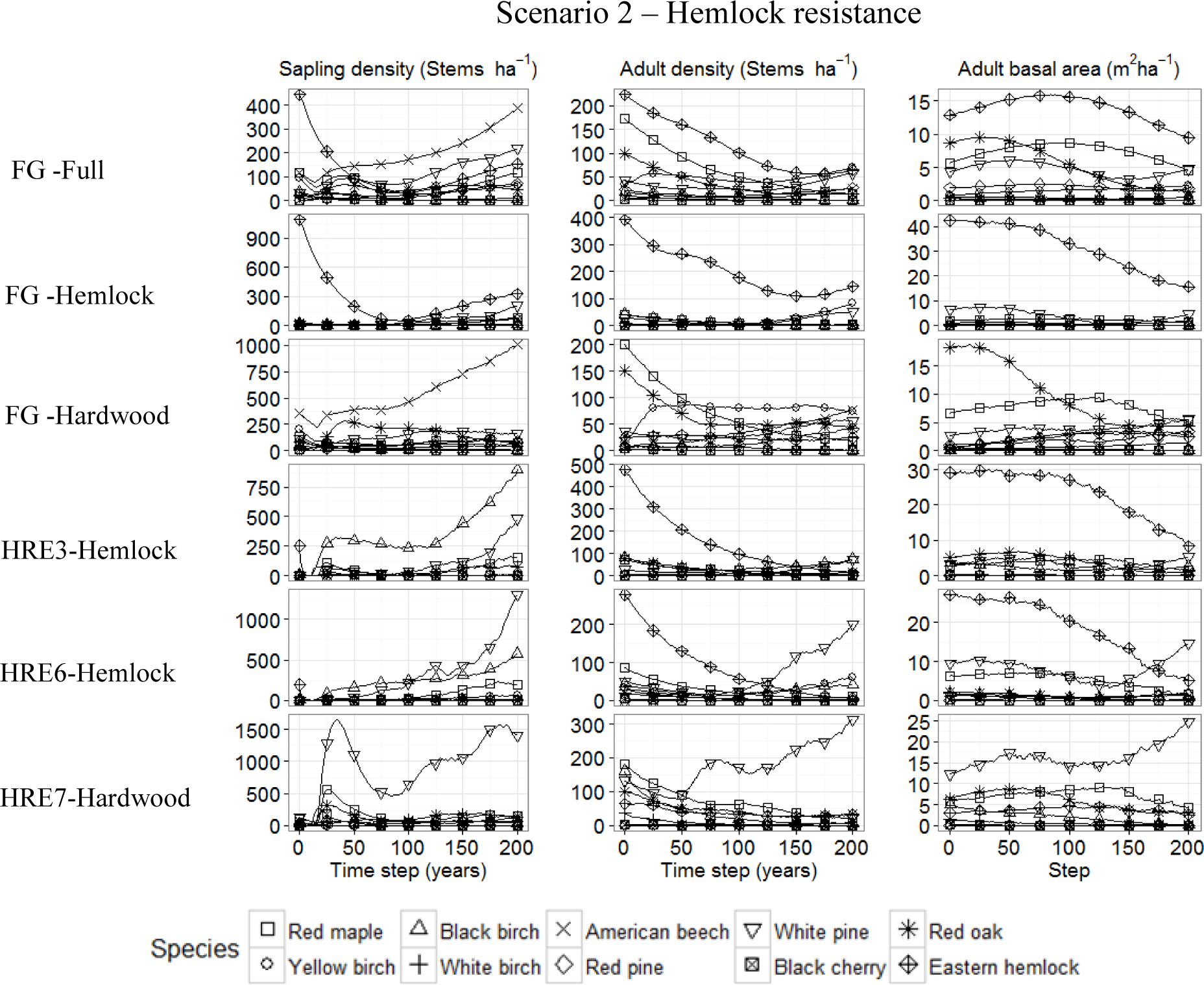
Simulated forest community dynamics over 200 years for Scenario 2 within the six study plots, characterized in terms of changes in sapling and adult stem density (stems ha^−1^) and adult basal area (m^2^ ha^−1^) for each of ten dominant tree species. Lines represent median values for each species of ten model simulations.

**Figure 8.**
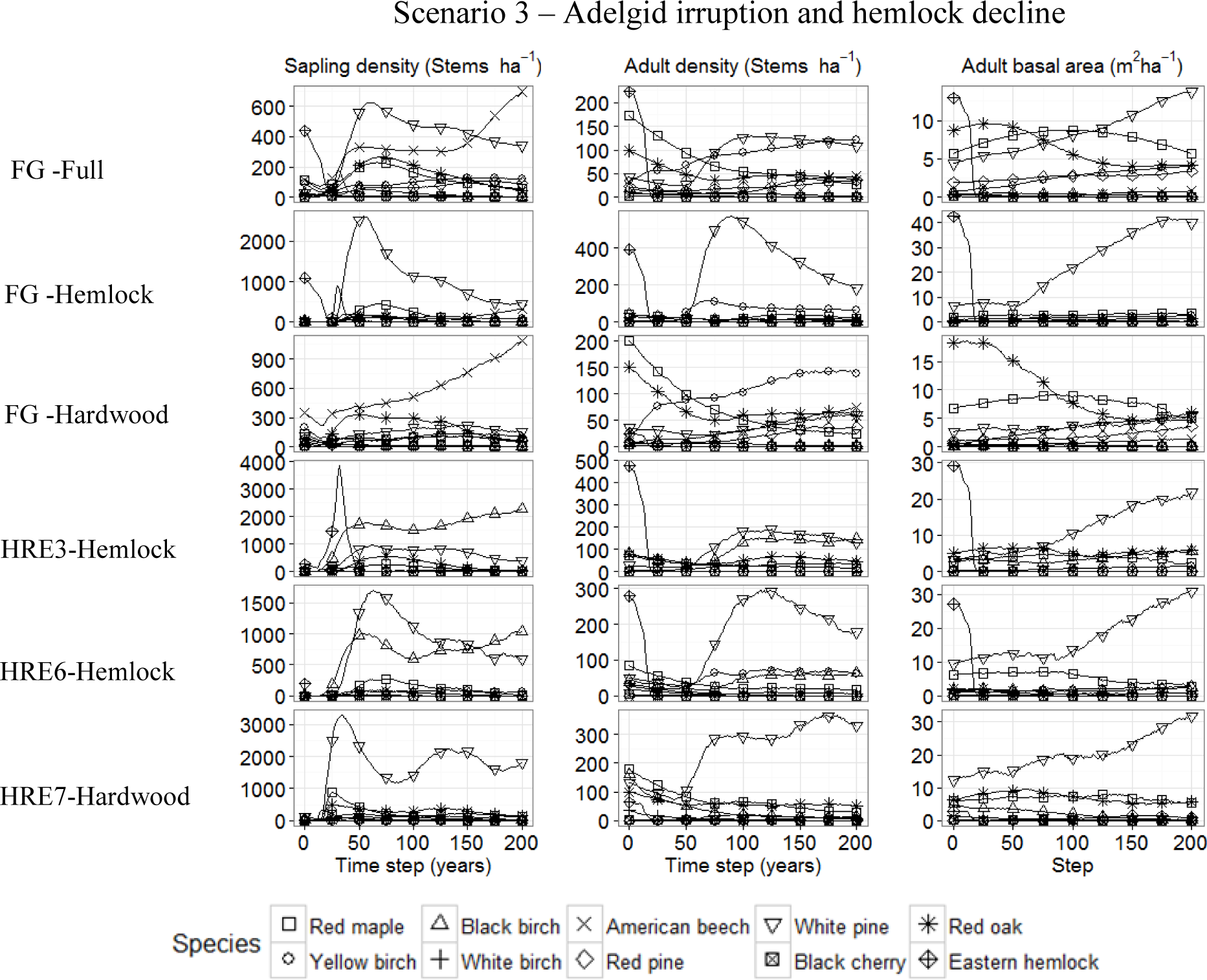
Simulated forest community dynamics over 200 years for Scenario 3 within the six study plots, characterized in terms of changes in sapling and adult stem density (stems ha^−1^) and adult basal area (m^2^ ha^−1^) for each of ten dominant tree species. Lines represent median values for each species of 10 model simulations.

**Figure 9.**
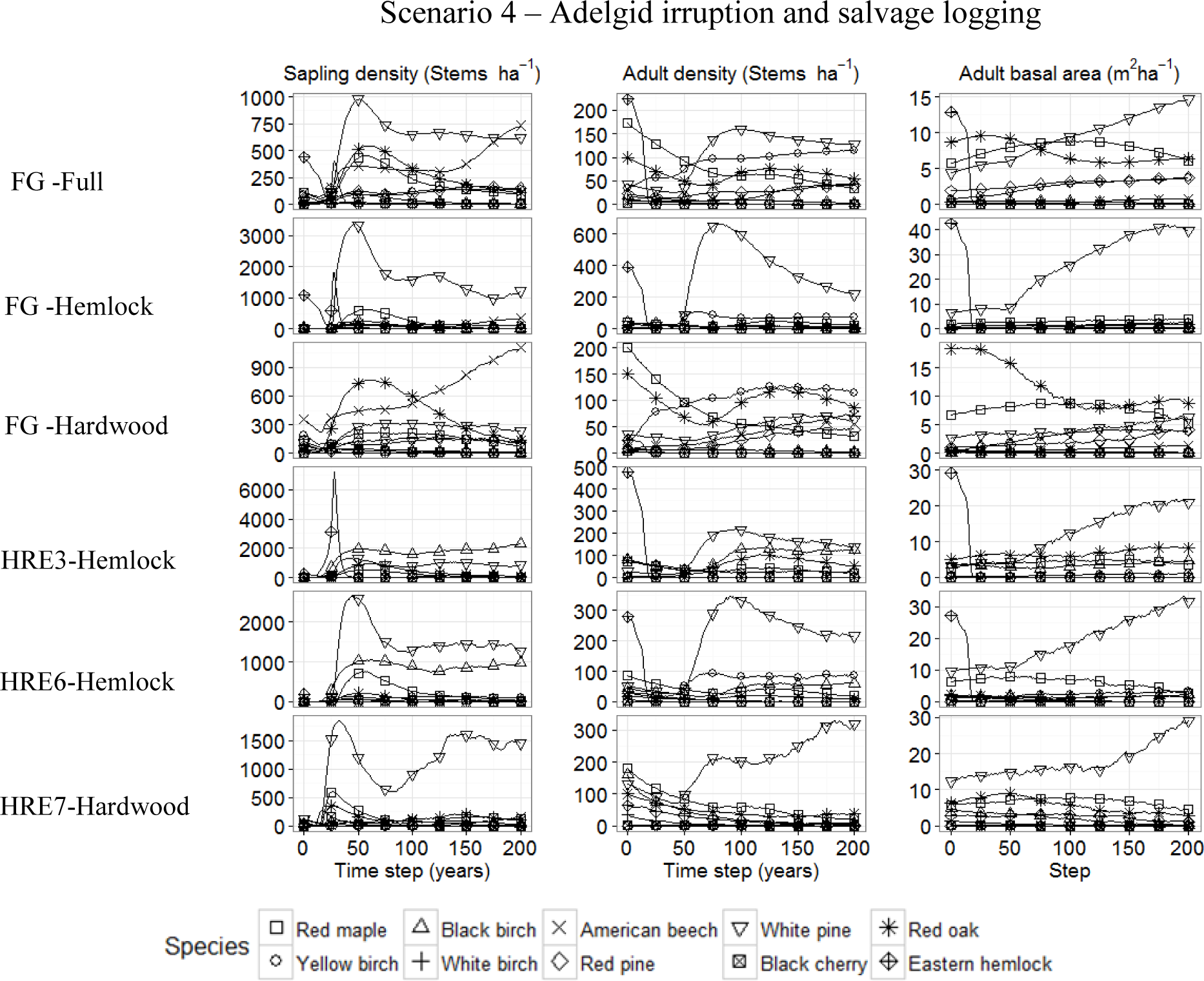
Simulated forest community dynamics over 200 years for Scenario 4 within the six study plots, characterized in terms of changes in sapling and adult stem density (stems ha^−1^) and adult basal area (m^2^ ha^−1^) for each of ten dominant tree species. Lines represent median values for each species of 10 model simulations.

SORTIE projected relatively similar 200-year trajectories under the three scenarios (Figs. 8 to 10) in which there was complete hemlock mortality by adelgids and natural snag dynamics (Scenario 3), “salvage logging” of hemlock just prior to death (Scenario 4), or preemptive logging of hemlock trees (Scenario 5). These scenario simulations suggested that, with exception of the HF-Hardwood plot, white pine is expected to increase significantly in abundance and basal area and will dominate these plots by year 200. In the FG-Hardwood plot, the loss of adult hemlock trees enabled American beech to recruit into the sapling tier canopy and yellow birch to become more prevalent in the adult tier. In both the HeRE6 and HeRE6 hemlock plots in these three scenarios, black birch also increased considerably in abundance over the 200-year period because of hemlock mortality. In year 200, total basal areas for these two scenarios (Fig. 6) were generally comparable to initial conditions, although in almost all plots there was a shift in structure to increased prevalence of larger trees, presumably due to the increase in large white pine canopy trees. The year-200 size structure in the HF-Hardwood plot, by contrast, contained fewer large trees, likely due to the large numbers of American beech recruits into this plot, which formed a dense canopy of sapling-sized trees.

**Figure 10.**
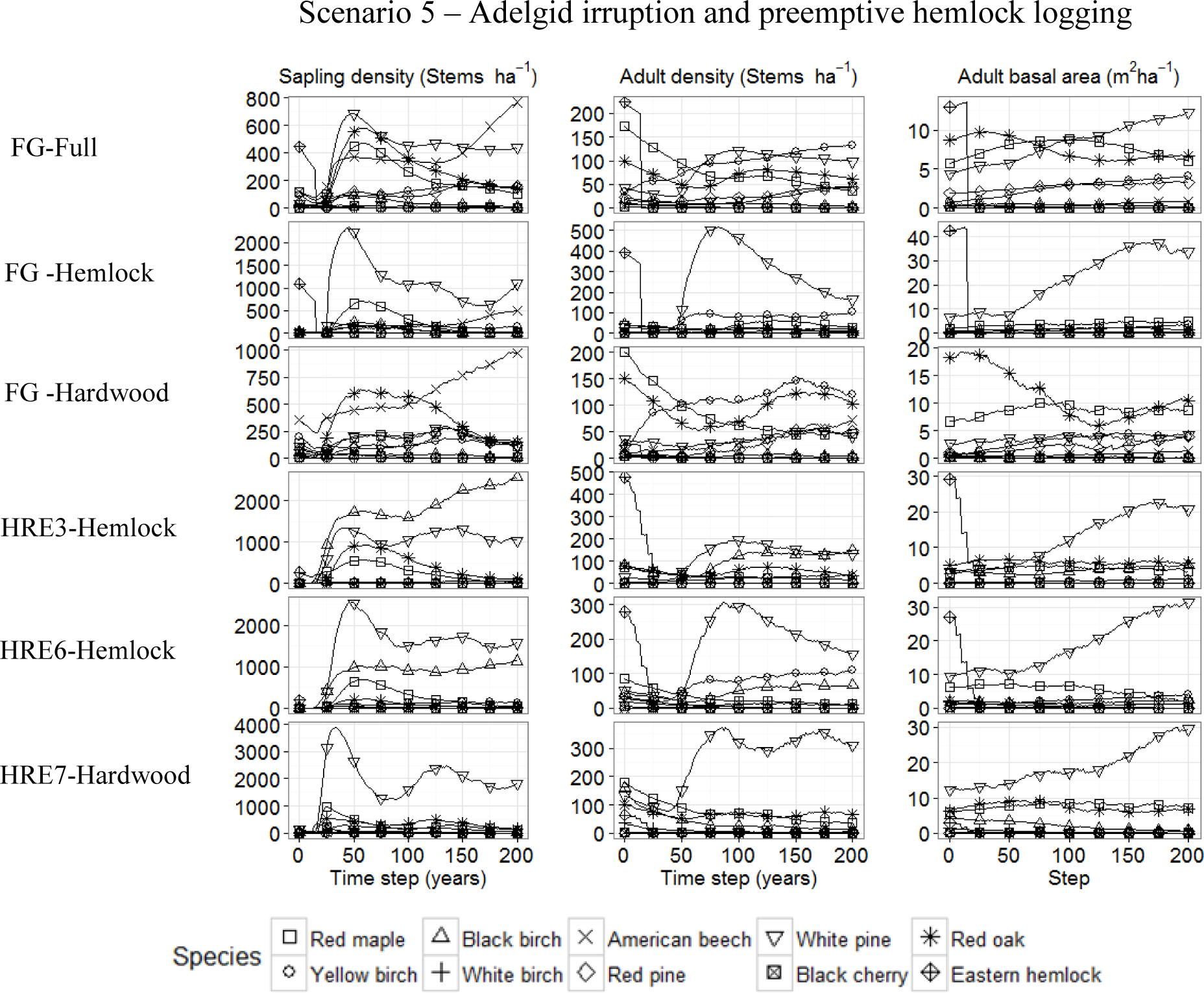
Simulated forest community dynamics over 200 years for Scenario 5 within the six study plots, characterized in terms of changes in sapling and adult stem density (stems ha^−1^) and adult basal area (m^2^ ha^−1^) for each of ten dominant tree species. Lines represent median values for each species of 10 model simulations.

Under a scenario of both hemlock and merchantable pine logging by year 25 (Scenario 6), differences in simulated community trajectories among the six plots depended on which species took most advantage of the increased light created by tree removal, and whether or not white pine was able to recover and gain dominance over time (Fig. 11). Across the full FG plot and within the FG-Hardwood subplot, American beech and yellow birch increased considerably in abundance in the sapling and adult tiers, respectively, preventing white pine from recovering in these plots. In the two HeRE hemlock plots (3 and 6), black birch became the most abundant species, although the white pine individuals remaining after logging were able to grow and become dominant in terms of basal area by the end of the simulation.

**Figure 11.**
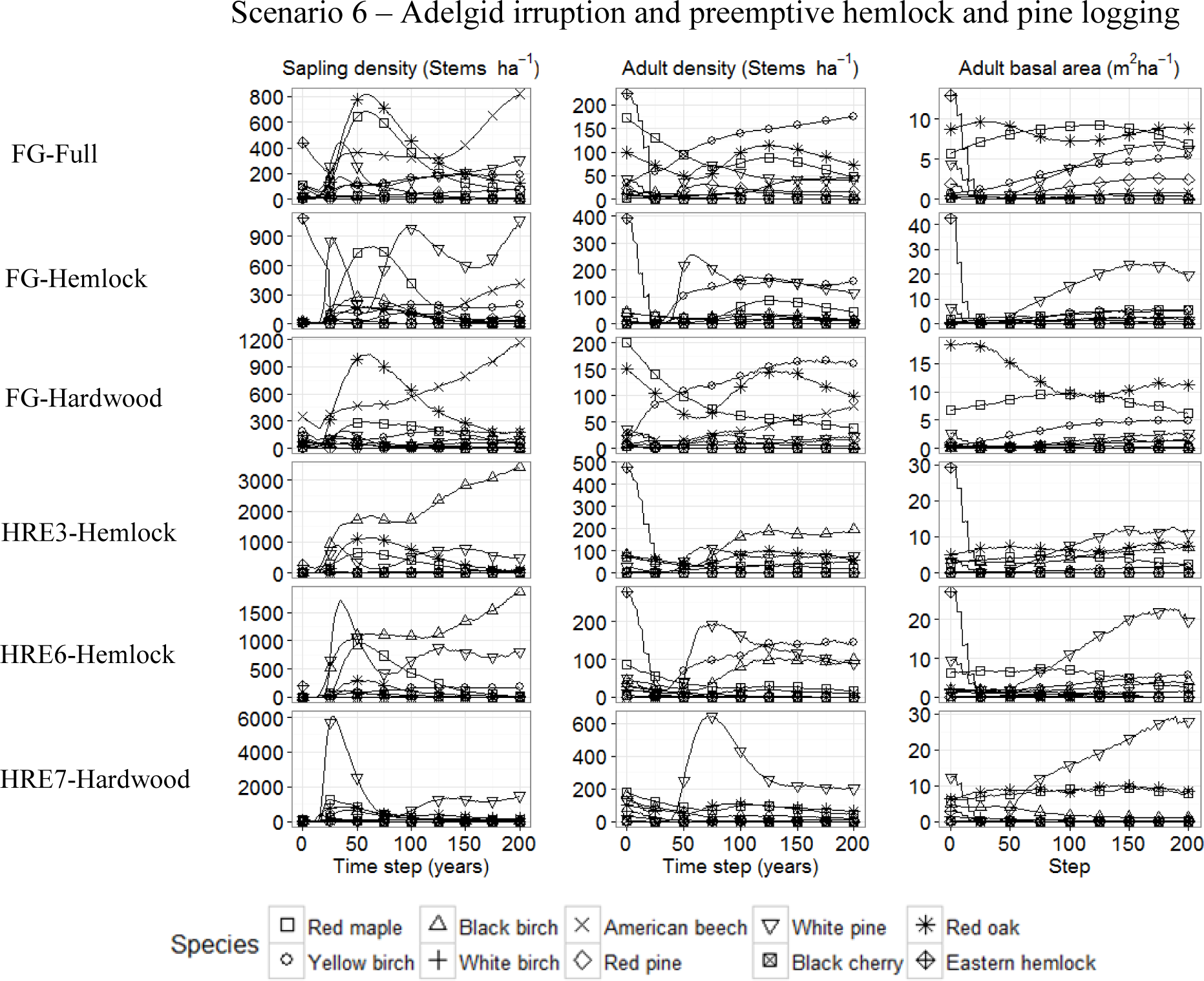
Simulated forest community dynamics over 200 years for Scenario 6 within the six study plots, characterized in terms of changes in sapling and adult stem density (stems ha^−1^) and adult basal area (m^2^ ha^−1^) for each of ten dominant tree species. Lines represent median values for each species of 10 model simulations.

### Forecasts – changes in diameter growth rates upon canopy removal

We examined projected species’ growth rates before (year 10) and shortly after (year 17) preemptive hemlock logging in Scenario 5 simulations to determine how the different species would respond to major increases in light and space after disturbance. These results indicated that all species would experience an increase in diameter growth upon the creation of canopy openings, at all growth stages (Fig. 12). However, growth rates, and the relative increases in growth after hemlock removal, differed significantly by species and study plot. White pine and red oak had the highest adult and sapling growth rates at the two times and across all plots; black birch also showed noticeable increases in year-17 growth rates in the two HRE-hemlock plots.

**Figure 12.**
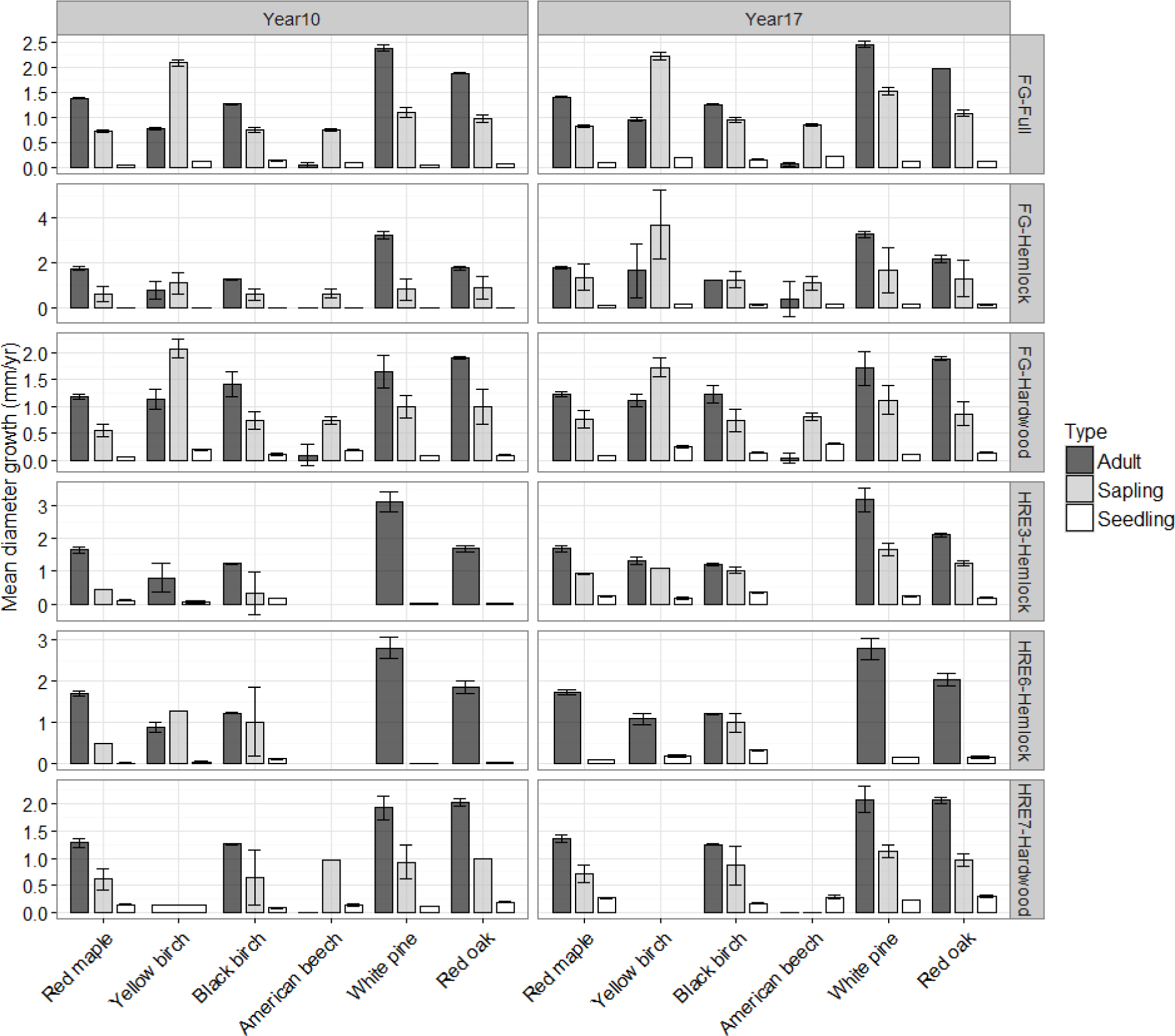
Mean annual diameter growth rates (mm year^−1^ ± 2 S.E.) for adults, saplings and seedlings in the six study plots, by species, based on Scenario 5 simulations. The left panel presents growth rates for simulation year 10, before preemptive harvesting of hemlock, and the right panel growth rates for year 17, two years after harvesting.

## DISCUSSION

There are gaps in our understanding about how forests will respond to the decline of particular tree species. Addressing such knowledge gaps in advance of the loss of foundation species, such as eastern hemlock, is particularly important because of the disproportionate effects foundation species have on the overall structure, composition, successional trajectory, and dynamics of the forest ecosystems they define. Given the long life spans of tree species and associated challenges with field experimentation, simulation modeling affords an essential tool for predicting how current forest conditions and core community dynamics processes may generate a range of new spatiotemporal realities for forests undergoing species decline and loss. Such modeling exercises also can inform the exploration and development of management strategies for at-risk forests. In this study, simulations with the SORTIE-ND individual-based model highlighted possible trajectories of secondary succession in these forests following adelgid-related hemlock mortality. Specifically, our modeling provides useful insights into how fine-scale, community dynamics may play out across a range of disturbance and initial condition scenarios.

Eastern white pine emerged in our simulations as the species able to take the greatest advantage of the increases of space and light created by the death of hemlock trees. Although generally not considered a highly versatile species, white pine is relatively shade tolerant, is a prolific and widespread disperser, has a broad ecological niche, and can be competitive in in a range of post-disturbance conditions (Abrams 2001). Indeed, white pine growth rates in our study plots in the years after hemlock death were high relative to the other species, making it highly competitive in our scenarios of hemlock decline, particularly in plots where there was a strong initial white pine adult component. In our study area within central Massachusetts, white pine and hemlock have had a long association in post-settlement forests, with hemlock dominating during more stable periods, and white pine taking advantage of gaps formed by periodic disturbances such as storms (Ireland et al. 2008). With increasing adelgid prevalence in this area, simulations suggest that white pine may replace hemlock as the dominant species, at least in the near term, in this novel disturbance scenario. Although some forest stands throughout New England have shown a general shift to a dominance of oaks and mixed hardwoods after significant hemlock dieback (Small et al 2005), our results indicate that even in our more hardwood-dominated plots, white pine will increase as the oak and red maple components of these stands mature and decline naturally over time.

Simulations also suggest that black birch would be competitive with white pine in hemlock gaps, but that its eventual relative abundance depends on some threshold of initial abundance and adequate seed production, as well as enough hemlock removed to provide ample light conditions for this relatively light-demanding species. This was particularly evident in the HeRE3 and HeRE6 hemlock plots, where black birch seedlings had relatively higher growth rates after hemlock harvesting than white pine (e.g., Fig. 12), but where slight differences in initial relative abundances and canopy gap sizes in these plots likely mediated the ability for black birch to take advantage of these better growth rates. In other locations in northeastern forests, where hemlock trees have died naturally due to adelgid impacts (Orwig and Foster 1998, Orwig et al. 2002), were killed experimentally (Orwig et al. 2013), or as a result of logging (Zukswert et al. 2014), abundances of black birch seedlings and saplings have also been observed to increase dramatically. Our results also suggest that, barring any other major disturbances, black birch could remain relatively dominant over the 200-year timespan in plots where it can establish and overtop other species. Black birch also survives several years in the seedbank in much greater numbers than other species (e.g. Farnsworth et al., Sullivan and Ellison). Our parameterization of SORTIE did not account for this effect, but it could favor establishment and subsequent dominance of black birch in the future.

SORTIE also projected a considerable influx and domination of the sapling tier by American beech upon the loss of hemlock in plots where there is sufficient initial occurrence of beech. Even in our hardwood-dominated plots, where there was relatively little hemlock present, the loss of hemlock individuals enabled beech to replace hemlock as the new shade tolerant species in the understory. Our adult growth parameterization for this species in SORTIE, based on previous FIA plot analyses by Canham et al. (2006, 2014), reflects a strong reduction in growth in beech beyond 15-cm DBH due to the effects of beech-bark disease over the past century. Indeed, beech-bark disease has been shown to cause a significant dieback of older trees and an increase in the suckering and abundance of sapling-sized beech, inhibiting the establishment of other, more common tree species; this scenario is playing out in many areas in the northeast USA (Garnas et al. 2011, Giencke et al. 2014) and is consistent with our simulation results. Such “beech brush thicket” conditions (Giencke et al. 2014) could persist indefinitely and define a new future reality for some stands in our study areas over the longer term.

On the whole, our results indicate that community trajectories and future forest structure will be qualitatively similar whether hemlock dies slowly from the adelgid, or is removed rapidly through logging. These results parallel findings of the Hemlock Removal Experiment, especially in comparisons between girdled and logging treatments (Lustenhouwer et al. 2012, Orwig et al. 2013, Ellison et al. 2014). Further, our “partial adelgid resistance” simulations (Scenario 2) indicate that despite significant initial (50-year) hemlock loss, its ability to cast such deep shade means that it can still retain relative dominance for a long period of time, preventing other species from taking quick advantage of new canopy openings while losing basal area itself. The result of this is effectively the creation of a “basal area suppression” situation, where hemlock persists but prevents other species from increasing in basal area compared to starting conditions.

A number of management options for hemlock forests undergoing decline, and the potential future implications of these options, have been explored previously (Brooks 2004, Orwig and Kittredge 2005, Foster and Orwig 2006, Fajvan 2008). These management options range from “doing nothing,” to carrying out silvicultural treatments or extensive hemlock logging. The first consideration arising from how to manage adelgid-affected hemlock forests relates to whether the main objective of such activities is the maximization of short-term profit or the long-term sustainability of the system. A focus on profit has led to the harvesting of hemlock and associated merchantable timber in advance of adelgid-induced mortality. Arguments against pre-emptive salvage logging include: (1) hemlock stands, even declining ones, form a unique and important ecosystem; (2) it is still as yet unclear whether there are, or will be, genetically-resistant hemlock individuals that could provide a refuge for future recovery or whether useful biocontrol measures may be developed in future years; (3) the responses of forest stands to levels of hemlock removal are site-specific, highly-variable, and therefore not always predictable. Our modeling highlights the last: the observed range of successional trajectories are contingent on local forest structure and dynamics. Thus, careful thought and additional research and modeling are needed to further explore future realities and whether human interventions of different types or intensities would produce a forest with compositional and structural conditions associated with sustainable ecosystem functions.

## ACKNOWLEDGEMENTS

We acknowledge support provided to BSC and HLB as part of the Charles Bullard Fellowship research program at Harvard University and Lincoln University. The Harvard Forest Hemlock Removal Experiment was established by, and is maintained as part of, the Harvard Forest LTER site, supported by the US National Science Foundation awards 0620443 and 1237491. The Harvard Forest CTFS-ForestGEO plot is also a project of the Harvard Forest LTER site. We offer heartfelt thanks to Charlie Canham and Lora Murphy, who have provided considerable expertise and data as input into the SORTIE-ND parameterization for our study sites.

